# Energy landscapes and heat capacity signatures for peptides correlate with phase separation propensity

**DOI:** 10.1101/2023.05.05.539523

**Authors:** Nicy, Jerelle A Joseph, Rosana Collepardo-Guevara, David J. Wales

## Abstract

Phase separation plays an important role in the formation of membraneless compartments within the cell, and intrinsically disordered proteins with low-complexity sequences can drive this compartmentalisation. Various intermolecular forces, such as aromatic–aromatic and cation–aromatic interactions, promote phase separation. However, little is known about how the ability of proteins to phase separate under physiological conditions is encoded in their energy landscapes, and this is the focus of the present investigation. Our results provide a first glimpse into how the energy landscapes of minimal peptides that contain *π*–*π* and cation–*π* interactions differ from the peptides that lack amino acids with such interactions. The peaks in the heat capacity (C*_V_*) as a function of temperature report on alternative low-lying conformations that differ significantly in terms of their enthalpic and entropic contributions. The C*_V_* analysis and subsequent quantification of frustration of the energy landscape suggest that the interactions that promote phase separation leads to features (peaks or inflection points) at low temperatures in C*_V_*, more features may occur for peptides containing residues with better phase separation propensity and the energy landscape is more frustrated for such peptides. Overall, this work links the features in the underlying single-molecule potential energy landscapes to their collective phase separation behaviour, and identifies quantities (C*_V_* and frustration metric) that can be utilised in soft material design.

## I. INTRODUCTION

Biomolecular condensates are membraneless organelles within the cell that are thought to form via phase separation of proteins and nucleic acids (Banani et al., 2017; Boeynaems et al., 2019; Brangwynne et al., 2009; Mittag and Pappu, 2022). Intrinsically disordered proteins are found ubiquitously in naturally occurring phase separating proteins (Dignon et al., 2018; Harmon et al., 2017; Jonas and Izaurralde, 2013; Malinovska et al., 2013; Pak et al., 2016; uiroz and Chilkoti, 2015; Schmidt and Görlich, 2015; Schuster et al., 2020; Uversky et al., 2015). Mutational studies have shown that *π*–*π* (aromatic–aromatic) and cation–*π* (cation–aromatic) interactions promote biomolecular phase separation, specially those involving tyrosine (Y), phenylalanine (F), and arginine (R) (Brady et al., 2017; Bremer et al., 2022; Fisher and Elbaum-Garfinkle, 2020; Greig et al., 2020; Lin et al., 2017; Martin et al., 2020; Nott et al., 2015; Qamar et al., 2018; Wang et al., 2018). In addition it has been demonstrated that some residues act as ‘stickers’ and promote phase separation, while other residues known as ‘spacers’ favour the solubility of proteins (Harmon et al., 2017^,2^01^8^; Holehouse and Pappu, 2018a). While at first glance, some stickers may contain similar functional groups, they can be unequal contributors to biomolecular phase separation. For instance, Y is better than F, and R is better than lysine (K) in stabilising condensates (Brady et al., 2017; Bremer et al., 2022; Fisher and Elbaum-Garfinkle, 2020; Greig et al., 2020; Lin et al., 2017; Martin et al., 2020; Nott et al., 2015; Qamar et al., 2018; Wang et al., 2018). R may also modulate phase separation in a context-dependent manner (Bremer et al., 2022). Some of these observations raise an important question: what are the key features that characterise the underlying energy landscapes of phase-separating proteins? In this paper, we address this question by applying the energy landscape framework to peptides with different sequences encoding *π*–*π* and cation– *π* interactions that are known to promote phase separation of proteins yielding biomolecular condensates. The energy landscape framework allows us to explore the potential energy landscape of the peptides by performing geometry optimisation to identify local minima and transition states, and connecting them via steepest-descent pathways (Wales, 2004). The energy landscape approach provides a powerful tool to explain emergent observable properties in terms of the atomic interactions at a fundamental level.

Specifically, we performed a computational analysis of the potential energy landscape for various hexapeptide monomers modelled at the atomistic scale. We chose hexapeptides because the secondary structure of pentapeptides is context-dependent (Kabsch and Sander, 1984), and therefore, the hexapeptides may represent the minimal system useful for investigating the conformational properties of peptides as well as the intramolecular interactions between the amino acids within a peptide monomer. Following the stickers-and-spacers model (Harmon et al., 2017; Holehouse and Pappu, 2018b; Yang et al., 2019) the hexapeptides are chosen to contain two dipeptide stickers joined together by a glycine–glycine (GG) spacer (Abbas et al., 2021). Working with such minimal systems allows us to directly link the differences in the energy landscapes to specific interactions between amino acid pairs, and hence, reduce the impact of cooperative and competitive effects.

A key signature of the energy landscape of a molecule is its heat capacity (C*_V_*). In this contribution we exploit the capability to produce rapid analysis of the heat capacity, and assign the peaks to specific local minima with distinct intramolecular interactions. Measurement of the C*_V_*, as a function of temperature can be useful to gain better insight into the thermodynamic properties of biopolymers, using differential scanning calorimetry (Benzinger, 1971; Cooper, 2010; Poland, 2001, ^2^00^2^; Prabhu and Sharp, 2005). Specific heat measurements for peptides at low temperatures can be employed as a measure of the elasticity of the molecule (Finegold and Cude, 1972). Here, we calculate the C*_V_* of peptide monomers using the harmonic superposition approximation (HSA) (Wales, 2017), and we observe features (peaks or inflection points) at low temperatures for the hexapeptides with phase separation promoting residues. The low-temperature peaks arise from competing structural motifs for a relatively small number of low-lying local minima. Peaks can be assigned to competition between these minima using the temperature derivative of the occupation probability (Wales, 2017). The theory provides an exact decomposition of C*_V_* in terms of local minima within the same approximation, which reveals the important cases of interest, where the peaks arise from competition between a few low-energy conformations. We emphasise that peaks in C*_V_* are simply being used as a diagnostic of the structure in the underlying landscape. This structure is clear in the harmonic normal mode approximation to the partition function and C*_V_* ; a more accurate treatment is not required to achieve this diagnostic.

The degree of frustration (Bryngelson and Wolynes, 1987; Onuchic and Wolynes, 2004) of the potential energy landscape, quantified via a frustration metric (De Souza et al., 2017), reveals the persistence of high energy barriers separating low-lying minima. In other words, the frustration reflects the existence of competing configurations. The frustration in the landscape is caused by different low-lying potential energy minima separated by significant barriers. Here, we find that the potential energy landscape is more frustrated for the peptides that contain residues (Y/R) with a higher propensity for phase separation, compared to the residues with a lower phase separation propensity. This observation agrees with the finding that the potential energy landscapes for intrinsically disordered proteins are multi-funneled (Chebaro et al., 2015). However, the frustration metric (De Souza et al., 2017) alone is not sufficient to predict phase separation propensity. Overall, we observe that the peptides with residues that have high phase separation propensity have distinct peaks or inflection points at low temperatures (significantly below the melting temperature) in C*_V_* plots and more frustrated potential energy landscapes. These features in C*_V_* correspond to competing structures stabilised by alternative interactions (aromatic–aromatic or cation–aromatic), or where the residues are oriented differently. This analysis suggests that the calorimetric criterion is a necessary but not a sufficient condition for phase separation (Zhou et al., 1999). The frustration metric provides an additional diagnostic to compare the phase separation propensity of residues in sequences that already exhibit features in C*_V_* at low temperatures.

## II. METHODS

The workflow adopted during the current study is presented in Fig. 1 and summarised below. The peptide sequences are constructed using the stickers-and-spacers model (Holehouse and Pappu, 2018b), and the hexapeptides are modelled using the FF99IDPS (Case et al., 2005; Wang et al., 2014) force field (Step 1, Fig. 1). The FF19SB (Tian et al., 2020) force field is also tested for some of the peptides to ensure that the structures represented by C*_V_* features depend on the interations within the sequence and not on the force field (Supplementary Information). The potential energy landscape is then explored using basin-hopping parallel-tempering (Step 2, Fig. 1) (Li and Scheraga, 1987, ^1^98^8^; Wales and Doye, 1997). Discrete path sampling (Wales, 2002) is employed to find the connected pathways between local minima (Step 3, Fig. 1). The convergence of sampling is monitored via disconnectivity graphs (Becker and Karplus, 1997; Wales et al., 1998) and heat capacities. The C*_V_* analysis is performed using the harmonic superposition approximation (Step 4, Fig. 1) (Wales, 2017), and the frustration in the landscape is quantified via a frustration metric (De Souza et al., 2017).

**FIG. 1:**
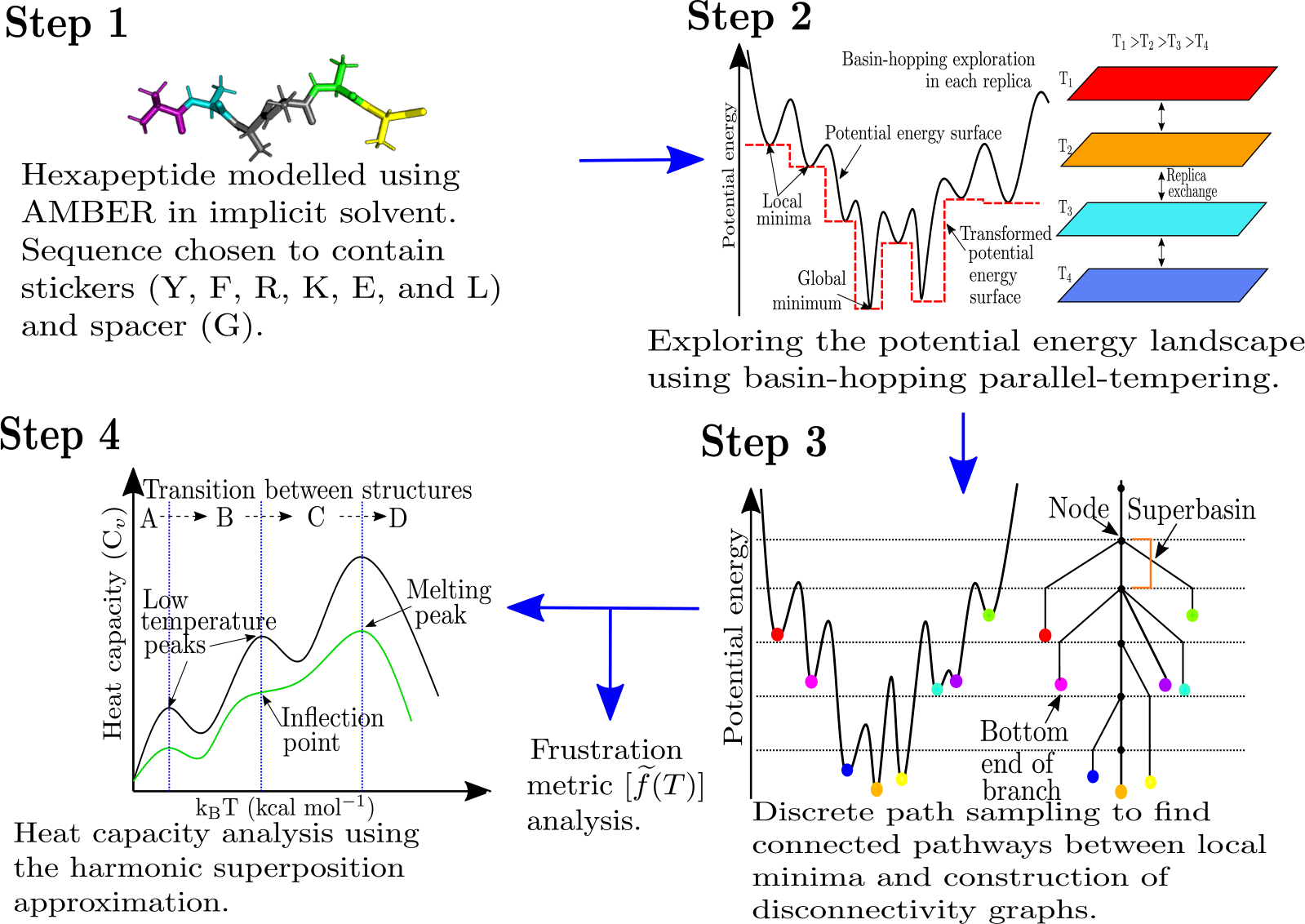
Schematic figure representing the workflow for the computational potential energy landscape exploration to interrogate peptides of varying phase separation propensities.

### Peptide model using AMBER

The hexapeptides are modelled using a properly symmetrised (Malolepsza et al., 2010) version of the FF99IDPs (Wang et al., 2014) force field along with an implicit solvent model (igb=8), and a monovalent ion concentration of 0.1 M (Case et al., 2005, ^2^02^2^). The N-and C-terminals are methylated, and methylamidated, respectively, to cap the charges in the zwitterionic form of the peptide (Step 1, Fig. 1). We also tested another force field FF19SB (Tian et al., 2020) and the uncapped peptides for both the force fields. The results are presented in the Supplementary Information.

### Basin-hopping parallel tempering

The global optimisation program GMIN (Wales, 2023a) is used to perform basin-hopping (Wales and Doye, 1997) parallel tempering (BHPT).(Li and Scheraga, 1987, ^1^98^8^) For the current computation, the AMBER interface with GMIN is used. A total of 16 replicas are used with temperatures exponentially distributed between 300 to 575 K. The exchanges are attempted at random with a mean frequency of 10, i.e., an average of one exchange after every 10 steps. The potential energy landscape is explored by performing 100,000 cartesian coordinate steps, and group rotation (Mochizuki et al., 2014) moves for the side chains. The local minima with C*_α_* in D- form and peptide bonds as cis-isomer are discarded. A root-mean-square (RMS) force convergence criterion of 10*^−^*^7^ kcal mol*^−^*^1^ Å *^−^*^1^ is employed to save the 400 lowest energy structures differing by at least 0.01 kcal mol*^−^*^1^ (to ensure uniqueness of local minima) after running BHPT (Step 2, Fig. 1).

### Discrete path sampling

Discrete path sampling (Wales, 2002) implemented in the OPTIM (Wales, 2023b), and PATH- SAMPLE (Wales, 2023c) programs is used to find optimal pathways between the local minima, and the global minimum. A discrete path is defined using elementary rearrangement between a local minimum, transition state, and another local minimum. The local minimum is defined as a stationary point with no negative Hessian eigenvalues whereas a transition state is a first-order saddle point with exactly one negative Hessian eigenvalue (Murrell and Laidler, 1968; Wales, 2004). The doubly-nudged (Trygubenko and Wales, 2004) elastic-band algorithm (Henkelman and Jónsson, 2000; Henkelman et al., 2000) is used to generate candidate transition states which are then refined accurately using hybrid eigenvector-following (Munro and Wales, 1999). Approximate steepest-descent is employed to find the local minima connected by the transition state using the limited-memory Broyden-Fletcher-Goldfarb-Shanno (L-BFGS) algorithm (Liu and Nocedal, 1989; Nocedal, 1980). Dijkstra’s shortest path algorithm (Dijkstra, 1959) is then used to choose the next pair of minima for which a new connection attempt is made, and the process is repeated until a fully connected pathway is found between the minima of interest using the missing connection algorithm (Carr et al., 2005). In the case of some peptides, chain crossing is observed. For these peptides, quasi-continuous interpolation (QCI) (Röder and Wales, 2018; Wales and Carr, 2012) is employed to find the correct pathways. Finally, the stationary point database is optimised using UNTRAP procedure (Strodel et al., 2007) in PATHSAMPLE to remove artificial frustration in the landscape; i.e., low-lying minima separated by large barriers where a lower energy transition state exists. The convergence of the stationary point database is monitored by the convergence of low-temperature peaks in C*_V_* plots, and by analysing the disconnectivity graph (Step 3, Fig. 1).

### Disconnectivity graphs

The potential energy landscape of a system of *N* atoms lies in a 3*N* + 1- dimensional space. Disconnectivity graphs provide a powerful way to visualise the multi-dimensional potential energy landscape (Becker and Karplus, 1997; Wales et al., 1998). They preserve the information about the minimum barrier for transitions between minima. The vertical axis of the disconnectivity graph represents the potential (or free) energy. The nodes on the vertical axis represent superbasins composed of disjoint sets of minima. Minima lying within the same superbasin can interconvert via a barrier less than or equal to the energy represented by the superbasin. Each branch originates from a node representing the superbasin and terminates at the energy of a local minimum corresponding to a single branch (Step 3, Fig. 1).

### Heat capacity analysis

The harmonic superposition approximation (which is accurate at low temperatures) can be used to express the total partition function as a sum of partition functions of all the local minima. The individual partition functions for the local minima are obtained using normal mode analysis, which yields the harmonic approximation to the vibrational density of states. The C*_V_* can now be expressed in terms of occupation probabilities of local minima and their temperature derivatives (Wales, 2017), i.e.,

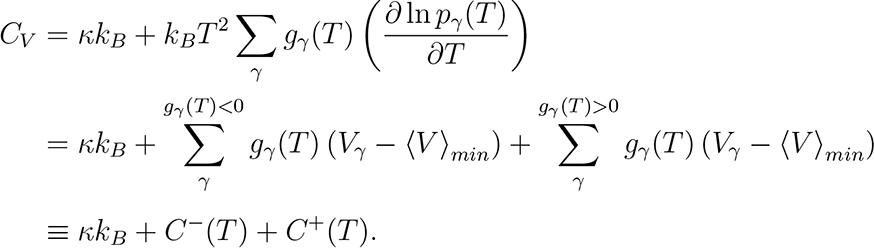

Here, *C_V_* is the heat capacity, *κ* = 3*N −* 6 is the number of vibrational degrees of freedom for a system of *N* atoms, *k_B_* is the Boltzmann constant, *g_α_*(*T*) *≡ ∂p_α_*(*T*)*/∂T* is the derivative of the occupation probability *p_α_* for minimum *α* with respect to temperature *T*, *V_α_* is the potential energy of minimum *α*, and *(V)_min_* is the mean potential energy of the minima. The peaks in C*_V_* represent transitions between states with decreasing (*g_γ_*(*T*) *<* 0) and increasing (*g_γ_*(*T*) *>* 0) occupation probability (Wales, 2017).

### Frustration metric calculation

Competing low-energy minima separated by significant barriers make the potential energy landscape frustrated. The frustration of the potential energy landscape can be quantified using a frustration metric (*f*^r^(*T*)), which is a function of temperature, i.e.,

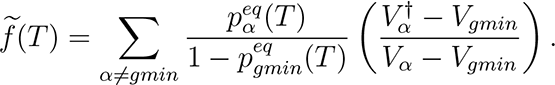

Here, *f* (*T*) is the frustration metric at temperature *T*, *V_gmin_* is the potential energy of the global minimum in the database, *V_α_* is the potential energy of minimum *α*, *V ^†^* is the potential energy of the highest energy transition state on the lowest energy pathway between *α* and global minimum, and *p^eq^*, and *p^eq^* are the equilibrium occupation probabilities of minimum *α* and global minimum which are calculated using the harmonic vibrational density of states. The global minimum does not contribute to frustration and its inclusion leads to erroneous decrease in frustration at low temperature. Hence the global minimum is excluded from the frustration metric calculation and occupation probabilities of remaining minima are renormalised (De Souza et al., 2017).

## III. RESULTS AND DISCUSSION

The importance of multivalency (Li et al., 2012), interaction strength (Asherie et al., 1996; Brangwynne et al., 2015; Choi et al., 2020; Das and Pappu, 2013; Hyman et al., 2014), and accessibility (Ruff et al., 2022) of stickers in promoting phase separation is well established. Here, we explore the energy landscapes (Fig. 2) of various hexapeptides containing a pair of dipeptides stickers separated by a GG spacer. The dipeptides stickers include FF, YY, RY, KY, YR, YK, RE, KE, FL, LF, and LL (Abbas et al., 2021). These sequences are chosen to encode the aromatic– aromatic, cation–aromatic, cation–anion and CH*· · · π* interactions. The interactions between individual pair of amino acids are further interrogated by analysing hexapeptides with a pair of stickers separated by two or four glycines. Energy landscapes are also explored for poly-amino acid hexapeptides containing a single type of amino acid residue including alanine (A), glycine (G), valine (V), arginine (R) and lysine (K). The peptide containing residues with better phase separation propensity show clear features in the C*_V_* at low temperatures (Fig. 3a, Section III A). These features are caused by competing low-energy conformations with different types of interactions (Fig. 4 and 5, Section III B). Further analysis of frustration of the energy landscapes reveals that the peptides with amino acids encoding better phase separation propensity result in more frustrated landscapes (Fig. 3b, Section III C). It is hypothesised that the collective behaviour of phase separation may be understood in terms of single-molecule properties by quantifying the heat capacity and frustration within the energy landscape framework. An interesting analog is how the existence of different conformations leads to polymorphic forms for various organic and inorganic molecules (Supplementary Information).

**FIG. 2:**
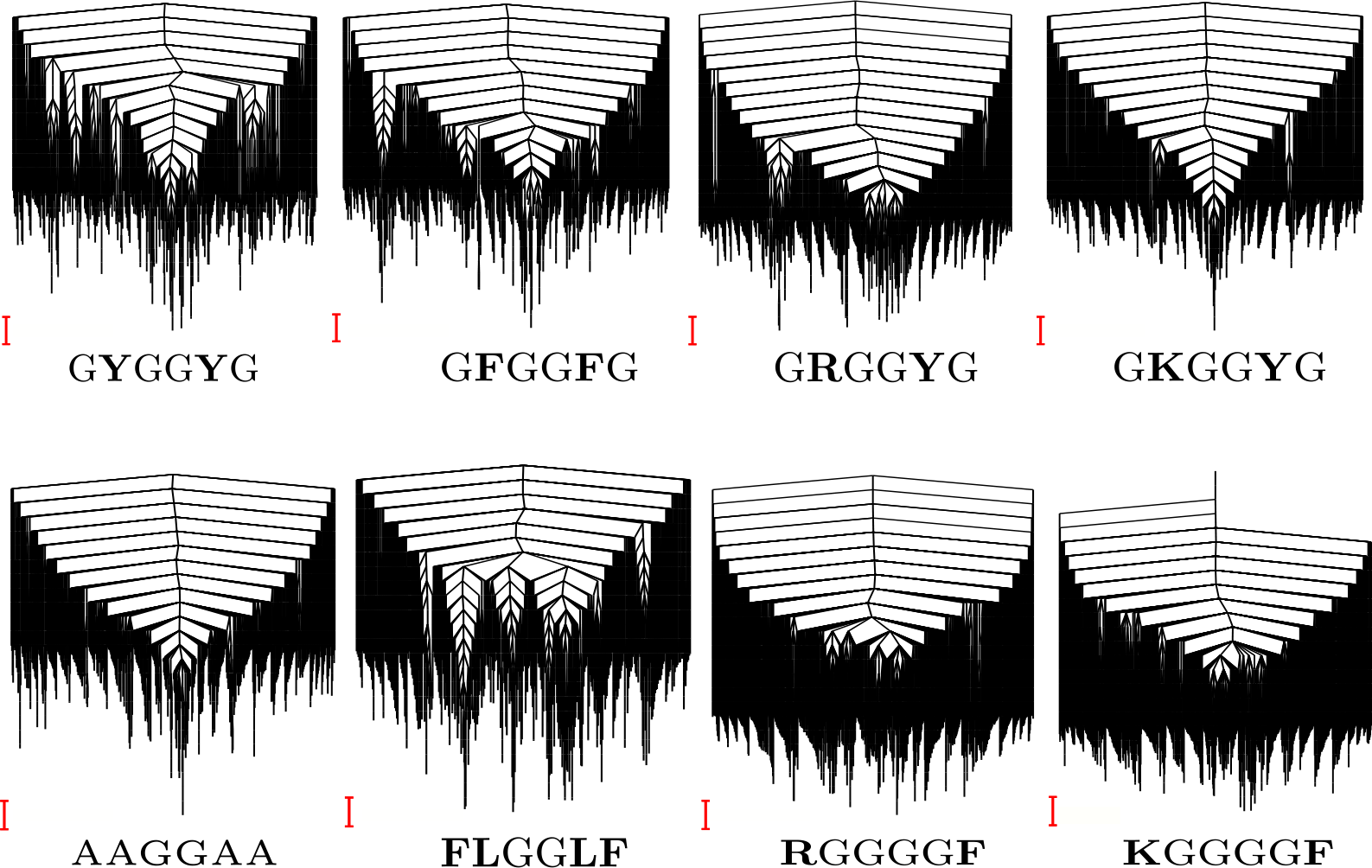
Representative disconnectivity graphs (Becker and Karplus, 1997; Wales et al., 1998) for some of the peptides studied. The scale bar is 1 kcal mol*^−^*^1^.

**FIG. 3:**
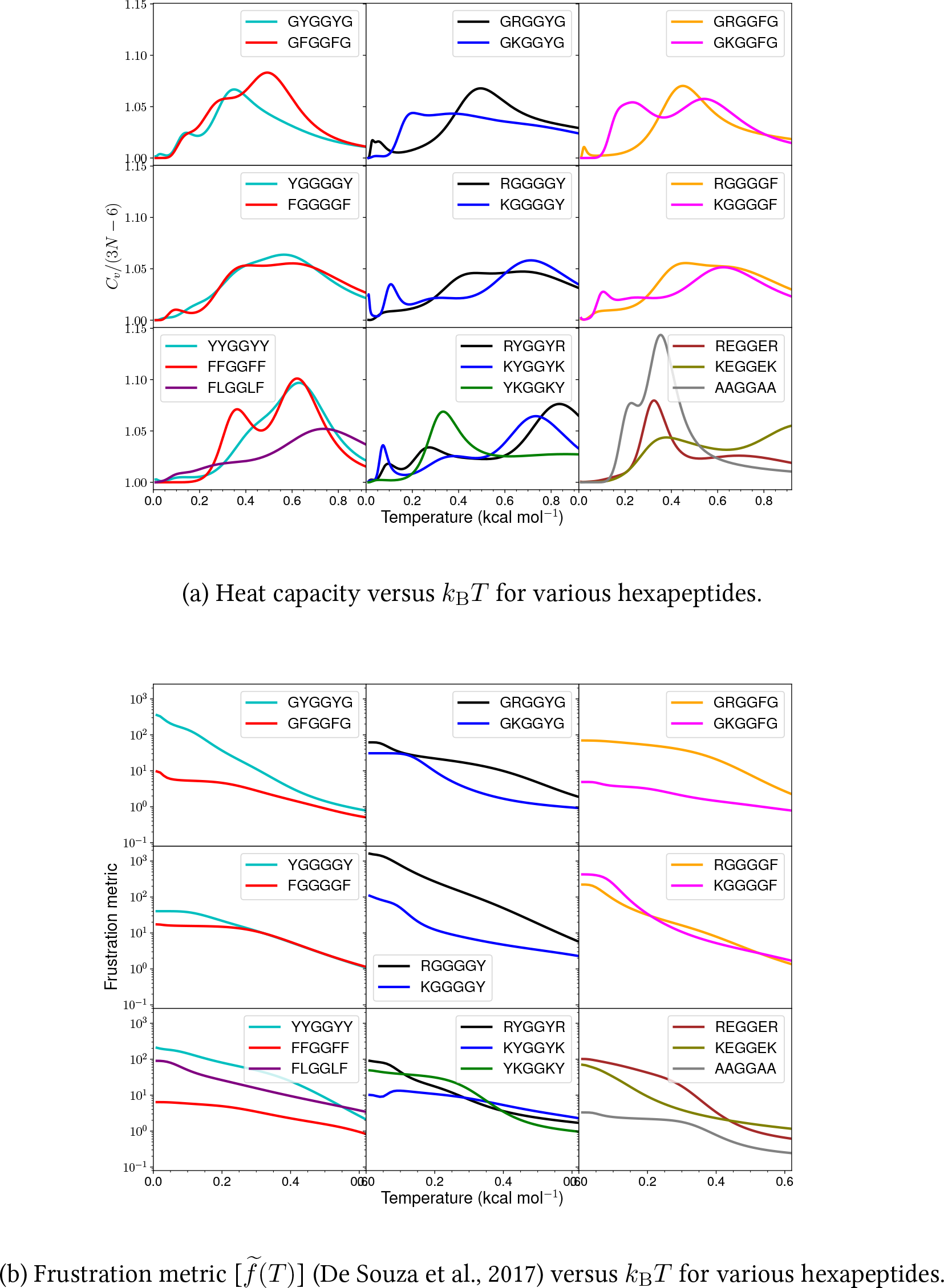
Heat capacity and frustration metric diagnostic for probing ph ase separation propensity encoded by different amino acid residues.

**FIG. 4:**
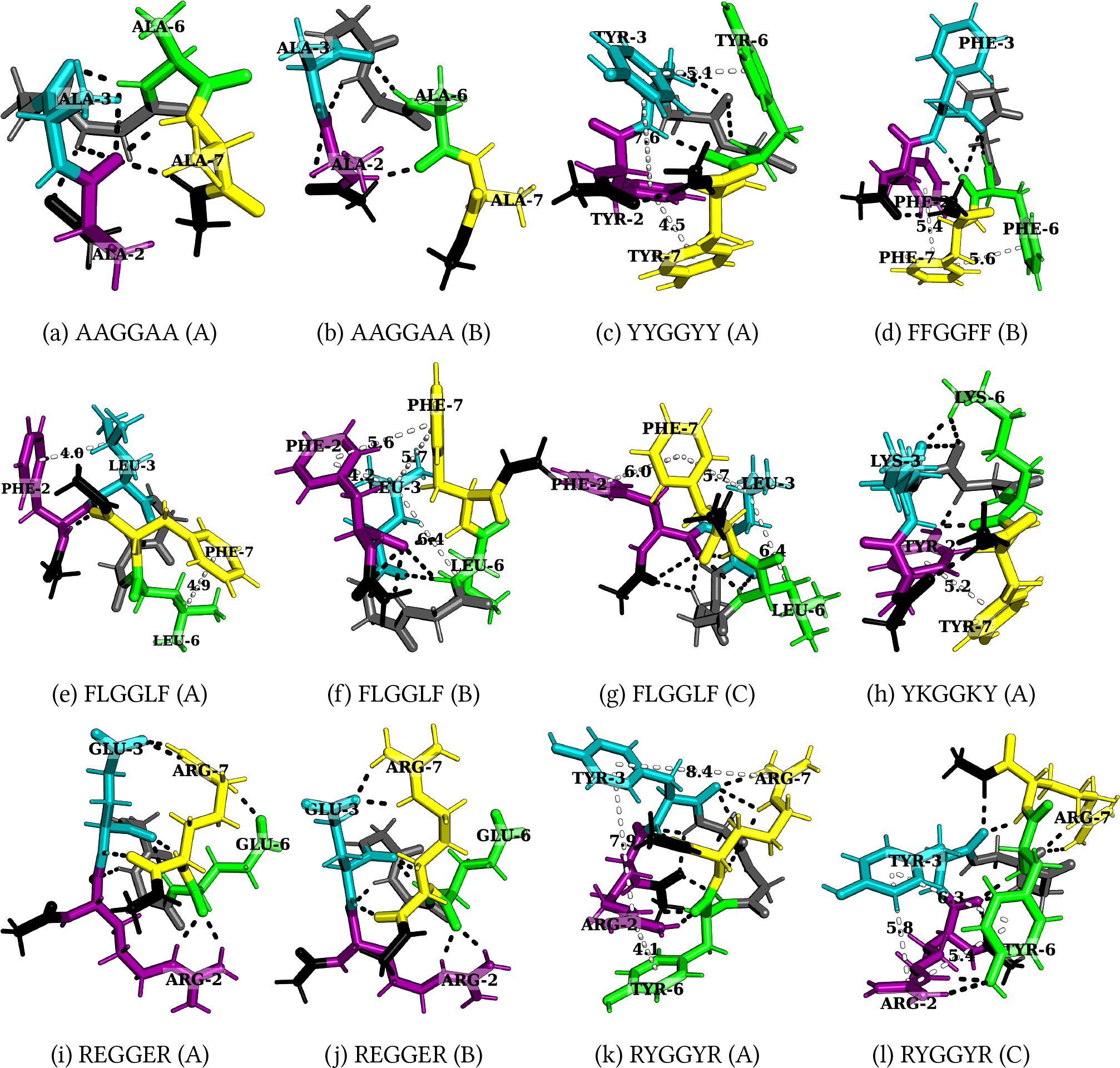
Structures corresponding to low-temperature heat capacity features. The first and second peaks correspond to the transition from A to B, and t hen from B to C, respectively.

**FIG. 5:**
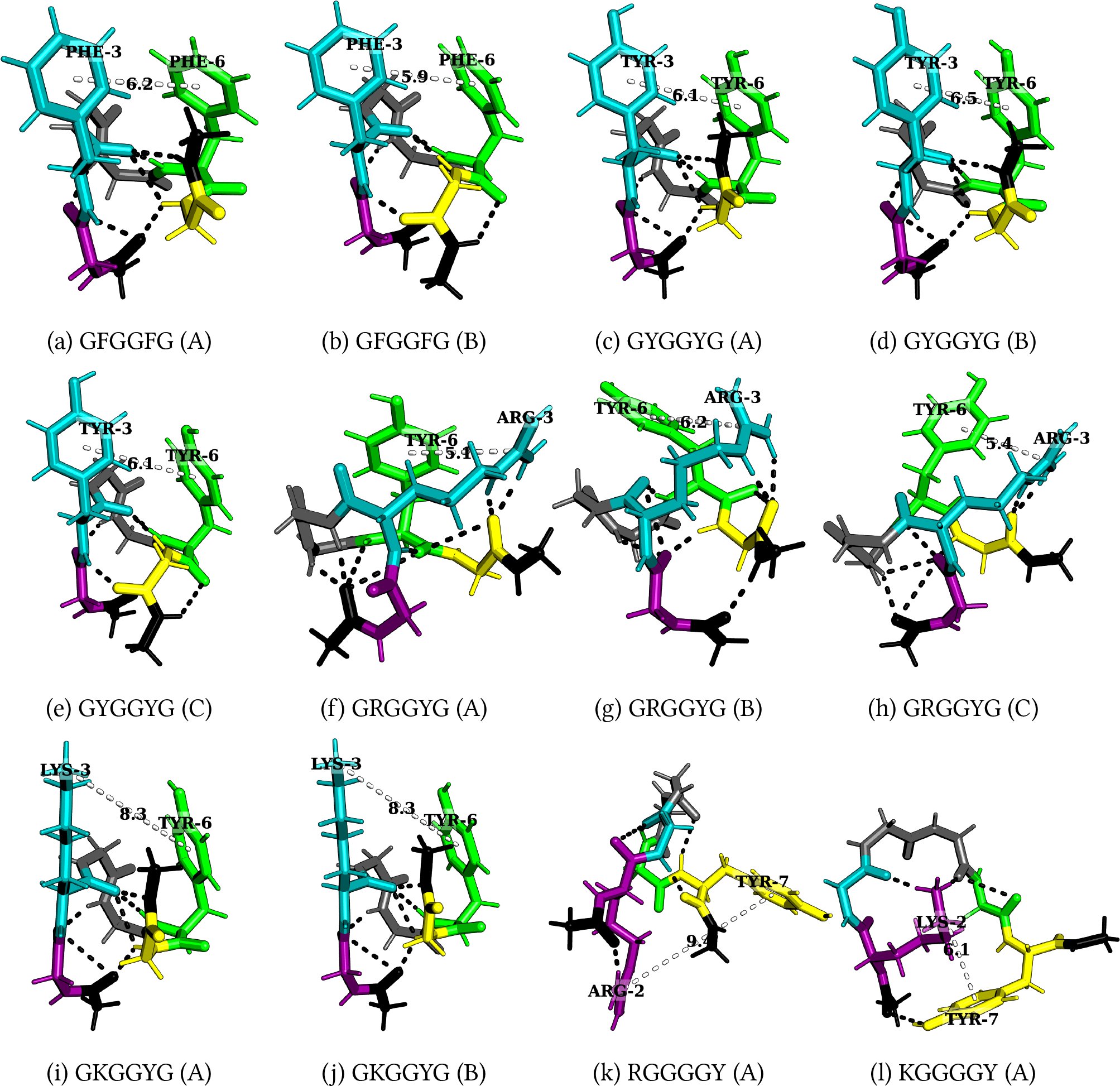
Structures corresponding to low-temperature heat capacity features. The first and second peaks correspond to the transition from A to B, and then from B to C, respectively.

### **A.** Heat capacity at low temperature

We first investigate the geometric, and energetic parameters that underlie the structural differences represented by low-temperature peaks in the heat capacity of peptides with varying phase separation propensities (Fig. 3a). We emphasise that we are using these features as a diagnostic for competing structures in the energy landscape, which may correlate with phase separation propensity. This computational construction does not need to be an accurate calculation of C*_V_*, nor does it need to be experimentally accessible. These peaks represent the transition between competing structures that have significant enthalpic and entropic differences and the integral over the peak represents the latent heat for this transition. In some C*_V_* plots, instead of distinct peaks, we observe distinct inflection points (GFGGFG, YGGGGY, RGGGGY, RGGGGF, YYGGYY, and FLGGLF) where the curvature of the plot changes. These inflection points (or shoulders) may be caused by overlapping peaks. The temperatures corresponding to these distinct inflection points are also considered, since they may also contain useful information. The hexapeptide AAGGAA is taken as the control, as it is predicted to have the lowest phase separation propensity of the set (Wang et al., 2018), and the C*_V_* is simpler (the potential energy landscape is not frustrated) compared to other peptides with more phase separation promoting residues. Note that it is not the height of the peaks but the existence of features at low temperature (below the melting temperature) that report on the structural heterogeneities in the landscapes and hence the phase separation propensities of the constituent residues in a sequence. Various other hexapeptides such as GGGGGG, AAAAAA, VVVVVV, EEEEEE, RRRRRR, and KKKKKK have also been analysed as controls and are found to show simpler C*_V_* profiles (Supplementary Information). However, distinct polar contacts between the main-chain atoms or between the main-chain, and side-chain atoms can produce features in C*_V_* (AAGGAA in Fig. 4a and 4b). In general, for hexapeptides with interactions that encode a higher propensity for phase separation, we observe more pronounced features (several distinct peaks and inflection points) in C*_V_* . The frustration metric can then be used as a further diagnostic. The C*_V_* plots for various other peptides are given in the Supplementary Information.

### **_B._** Interactions leading to features in C***_V_***

A low-temperature heat capacity peak often arises from a transition from a compact structure with two pairs of dominant interactions between four residues (YYGGYY - Fig. 4c and FLGGLF - Fig. 4e) to another structure with a similar pair of interactions, but with residues oriented differently, or a relatively extended structure with two pairs of dominant interactions between three residues (FFGGFF - Fig. 4d, and FLGGLF - Fig. 4f and 4g). Depending on the number of stickers in the peptide, the low-temperature peak may also correspond to a transition from two pairs of dominant interactions between three residues to one pair of interaction between two residues (KEGGEK, and REGGER - Fig. 4i and 4j). A detailed discussion of the competing structures for various hexapeptides is given below.

### Tyrosine vs phenylalanine

The presence of a hydroxyl group in tyrosine not only enhances its hydrogen-bonding ability, but also results in different rotamers leading to features in C*_V_* at low temperatures (GYGGYG, and YGGGGY). GFGGFG and GYGGYG exhibit inflection points, and distinct peaks at low temperatures, respectively. In particular, the low-temperature feature in GFGGFG and FGGGGF corresponds to the transition between a structure with methyl–aromatic and aromatic–aromatic interactions to a structure with an aromatic–aromatic interaction, which further changes to a structure with several polar contacts between distinct atoms. In contrast, for GYGGYG, and YGGGGY, the features at low temperature correspond to the transition between rotamers of the aromatic ring containing methyl–aromatic, and aromatic–aromatic interactions. Here, the methyl group belongs to the C-terminal cap of the peptide. Interestingly, the observation of a low-temperature peak resulting from the presence of ring rotamers can be compared to an experimental observation in which a bulge in the C*_V_* plot of polystyrene was attributed to the rotation of the phenyl ring around the chain axis (Warfield and Petree, 1962). The orientation that optimises the aromatic interaction depends on the distance between the C*_α_* atoms, stacked at a short distance, and T-shaped at a longer distance (Chelli et al., 2002; Hunter et al., 1991). Offset-stacked structures can also be energetically favourable (Ninković et al., 2014), and the methyl group of cap can also interact with an aromatic residue (Zanuy et al., 2004). We observe similar edge-to-face, C-H*· · · π*, and methyl–aromatic interactions for GFGGFG (Fig. 5a and 5b), GYGGYG (Fig. 5c, 5d, and 5e), FGGGGF, and YGGGGY.

### Arginine vs lysine

GRGGYG exhibits features in C*_V_* because of the interaction between R and Y, and the presence of ring rotamers (rotamer of an aromatic ring) for Y (Fig. 5f, 5g, and 5h), whereas for GKGGYG, and GKGGFG, it is the methyl group in the C-terminal cap that preferably interacts with the Y/F (Fig. 5i and 5j). We still see features in C*_V_* for GKGGYG because of ring rotamers for Y. In the case of RYGGYR, one of the peaks corresponds to the structural transition between the aromatic–cation–aromatic interaction motif to the aromatic–cation interaction motif (Fig. 4k and 4l). Hence, R has more propensity than K to interact with the aromatic residues.

### Context-dependence

Phase separation may be regarded as a percolation network transition (Mittag and Pappu, 2022). The difference in size, the steric packing of R, and K, the number of spacers between the stickers, and the distance between the stickers may be useful in explaining the context-dependent properties of these amino acid residues in terms of accessibility and networking ability of stickers to interact with each other. Consider the peptides RYGGYR, GRGGYG, GKGGYG, RGGGGY, and KGGGGY. Even though the presence of R leads to more features in the C*_V_* plots, we observe that when the cationic and aromatic residues are far apart, as for RGGGGY, and KGGGGY, K seems to be more flexible, and less sterically inhibited, and therefore, it can interact well with Y, whereas R seems to be more rigid and does not interact favourably with Y/F (Fig. 5k and 5l). Previous reports suggest that K/RNA coacervates are more dynamic than R/RNA coacervates (Ukmar-Godec et al., 2019), and the R-rich motif may act as a phase disruptor (Odeh and Shorter, 2020). While the different behaviours of R and K may be understood in terms of the relative strength of the interactions, it is also possible that the flexible nature of K compared to R may play a role. Furthermore, the shuffling of sequence may alter the presence of charged residues near the N-/C-termini, which may lead to differences in the properties of these peptides because of the charge interaction with the peptide dipole. The dipole moment effect is expected to be more significant in the case of an uncapped peptide in zwitterionic form (Marqusee and Baldwin, 1987; Tkatchenko et al., 2011).

### Aromatic–aromatic vs cation–aromatic interactions

Favourable cation–aromatic interactions between R and Y are observed in RYGGYR (Fig. 4k and 4l). However for YKGGKY, the aromatic–aromatic interaction between two tyrosine residues is preferred over the cation– aromatic interaction between K and Y (Fig. 4h). This observation hints at the role played by the proximity of interacting residues in a sequence, i.e., the two tyrosine residues located at the ends can establish an aromatic-aromatic interaction, which is preferred over the weaker interaction offered by the lysine residues. From a broader perspective, this result may be useful in understanding the context-dependent properties of amino acid residues across different sequences.

### Cation–anion interaction

Hydrogen-bonding between oppositely charged amino acids may lead to the formation of salt bridges where the same residue interacts with two different residues (complex) or between two oppositely charged residues (simple) (Musafia et al., 1995). Both REGGER (Fig. 4i and 4j), and KEGGEK exhibit low-temperature C*_V_* peaks corresponding to the transition from structures containing a complex salt bridge to a simple salt bridge. The complex salt bridge is formed by the interaction of the same cationic residue with two anionic residues. The next C*_V_* peak at a higher temperature corresponds to the transition from a structure with a cation interacting with a particular anion to a structure with the same cation interacting with a different anion in a different orientation as in the case of uncapped KEGGEK peptide.

### Partial phase separation

Leucine and phenylalanine are constituents of peptides exhibiting partial phase separation (Abbas et al., 2021), and the C*_V_* plot for FLGGLF contains features at low temperatures. The peak represents the transition from a structure containing two distinct pairs of L-F interactions, arising from four residues, to a structure with two pairs of interactions arising from three residues F, F, and L (Fig. 4e, 4f, and 4g). Several C-H*· · · π* interactions can occur between L and F. Hence, partial phase separation may occur for peptides containing amino acids capable of exhibiting distinct pairs of interactions. However, the interaction strength between stickers is weaker compared to the cation/aromatic–aromatic interaction. Although weak, the C-H*· · · π* interaction is known to play an important role in supramolecular organisation (Piccolo, 2001).

### **C.** Frustration in energy landscape

The frustration (Bryngelson and Wolynes, 1987; Onuchic and Wolynes, 2004) in the multi- dimensional potential energy landscape can be visualised by analysing the multiple funnels in the disconnectivity graph representation (Becker and Karplus, 1997; Wales et al., 1998) (Fig. 2), i.e., more funnels with low energy minima separated by significant barriers from the global minimum make the landscape more frustrated at low temperatures. uantitatively, we can calculate a frustration metric (De Souza et al., 2017), which is generally larger for peptides containing Y/R than for peptides containing F/K at lower temperatures (Fig. 3b). In particular, at a very low temperature corresponding to *k_B_T* = 0.2 kcal mol*^−^*^1^, the frustration metric for GYGGYG is 8 times the value for GFGGFG, YYGGYY is 16 times larger than FFGGFF, GRGGYG is 2 times larger than GKGGYG, GRGGFG is 16 times larger than GKGGFG, and REGGER is 5 times larger than KEGGEK. At *k_B_T* = 0.1 kcal mol*^−^*^1^ the frustration metric of RYGGYR is 3 times greater than that of KYGGYK. Hence, it appears that the frustration in the landscape for the monomer peptide directly correlates with the relative phase separation propensity of its constituent residues. Interestingly, KGGGGF is 3 times more frustrated than RGGGGF at very low temperature (*k_B_T* = 0.1 kcal mol*^−^*^1^). Here, the larger number of spacers (4 glycines) increases the distance between the stickers and affects the inaccessibility. The accessibility is reduced more in the case of R, which appears more rigid compared to the more flexible K residue. This difference may explain the context-dependent properties of R in phase-separating proteins. Moreover, the potential energy landscape of FLGGLF is 5 times more frustrated than FFGGFF at a very low temperature (*k_B_T* = 0.2 kcal mol*^−^*^1^). However, FFGGFF has distinct peaks in the C*_V_*, in contrast to FLGGLF (Fig. 3a). These features are caused by a stronger aromatic–aromatic interaction between two F, which correlates with the better phase separation propensity of residues in FFGGFF, whereas the interaction between F and L may facilitate partial phase separation (Abbas et al., 2021). The frustration metric plots for various other peptides are given in the Supplementary Information.

## IV. CONCLUSIONS

We have investigated the hypothesis that the energy landscape of peptide monomers may report on their phase separation ability, which is a collective phenomenon. The different possible arrangements in which the aromatic–aromatic, and cation–aromatic interactions can occur in a peptide monomer can produce low-temperature peaks in the heat capacity. Additionally, the high barriers between the alternative low-lying potential energy minima and the existence of several such conformations, as visualised by multiple funnels in the disconnectivity graph, report upon the highly frustrated potential energy landscape of the system. Together, features in the heat capacity plot, and the frustration of the landscape, quantified using the frustration metric, appear to correlate with increased phase separation propensity of the constituent residues. This analysis also provides a useful framework to investigate the context-dependent properties of amino acid residues in different sequences. While there have been several attempts (Dzuricky et al., 2020; Simon et al., 2017) to guide the rational design of peptides useful for bioengineering applications, the present study presents a new perspective to design peptides with targetted phase separation behaviour. It is important to understand that we are not actually interested in the low-temperature behaviour of the heat capacity and that an accurate calculation is not required. Rather, we are using peaks in an approximate C*_V_* as a computational construction to diagnose competition between alternative favourable structures. It is the characteristics of these conformations that may provide a structural interpretation and diagnostic of higher-order behaviour in condensates such as liquid-liquid phase separation. Our results suggest that there may indeed be such a connection. We do not claim that this connection is universal, but we do suggest that it may be useful.

## AUTHOR CONTRIBUTIONS

J.A.J, R.C.G, D.J.W and Nicy conceived the idea and designed the study. Nicy performed the simulations and wrote the first draft. All the authors helped with the analysis, interpretation of data and corrected the final draft.

## DATA AVAILABILITY STATEMENT

The discrete path sampling databases can be obtained from authors upon request.

## SUPPLEMENTARY INFORMATION

Effect of capping peptides: The methyl group at the C-terminal cap may interact with aromatic residues (GFGGFG, and GYGGYG). The polar contacts established between caps may constrain the structure, and the loss of these polar contacts may help the stickers interact with each other better (GRGGFG). When the cap interacts with a sticker, the distance between the stickers may be larger to minimise the repulsive interaction between the cap and the other sticker. In contrast, when the cap is not interacting with the sticker, it may be possible for the stickers to establish better contact by reducing the distance between them (GFGGFG, GYGGYG, GKGGFG).

Weak interactions in polymorphism Several weak interactions play an important role in driving phase separation, such as quadrupole-quadrupole (Burley and Petsko, 1985), charge- quadrupole interactions (Burley and Petsko, 1986), and hydrogen-bonding (Burley and Petsko, 1988). These weaker interactions are anisotropic, and for the same pair of residues these can be alternative favourable conformations, which may then pack into different arrangements (Berkovitch-Yellin and Leiserowitz, 1984), possibly leading to the existence of different phases. This suggestion is consistent with the conformational polymorphism (Nangia, 2008) seen in organic crystals. While the distinction between competing intramolecular and intermolecular interactions is beyond the scope of the current study, the smaller and larger distances between the interacting stickers within the monomers may be useful in visualising the various possible ways in which the interactions may occur in oligomeric systems. Polymorphism is also known to occur in amino acids (Purohit and Venugopalan, 2009), and protein complexes (Tompa and Fuxreiter, 2008), and H-bonding ability is important in polymorphic molecules (Cruz-Cabeza et al., 2015; Kumar and Nangia, 2014). The stability of non-linear H-bonds may not differ significantly from the collinear H-bonds (Baker and Hubbard, 1984). Metastable conformers may be stabilised by favourable packing arrangements (Babu et al., 2010), and some polymorphs are known to have different strengths for *π*–*π* interactions (Fan et al., 2009).

**FIG. 6:**
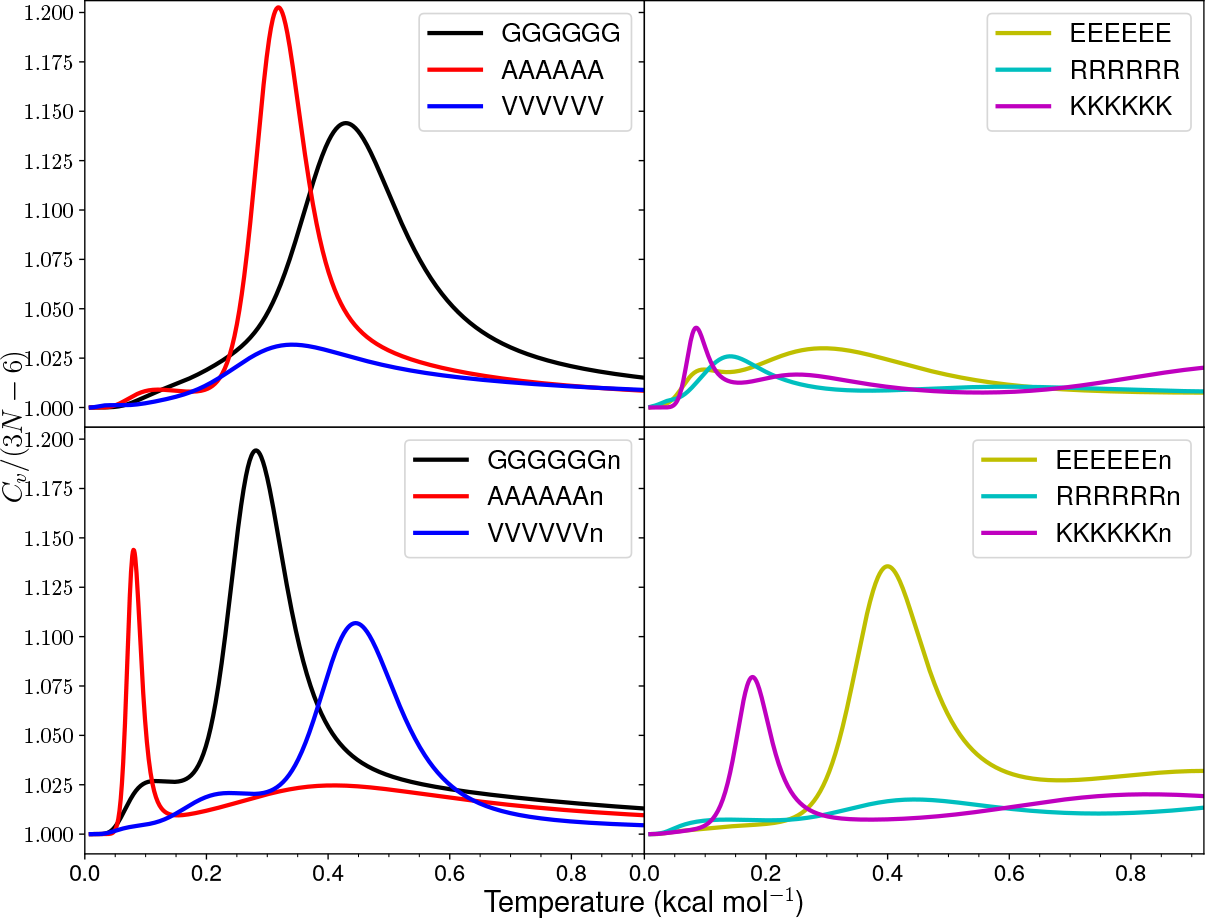
Heat capacity versus *k*_B_*T* for various hexapeptides. The uncapped peptide sequences end with a lowercase ‘n’.

**FIG. 7:**
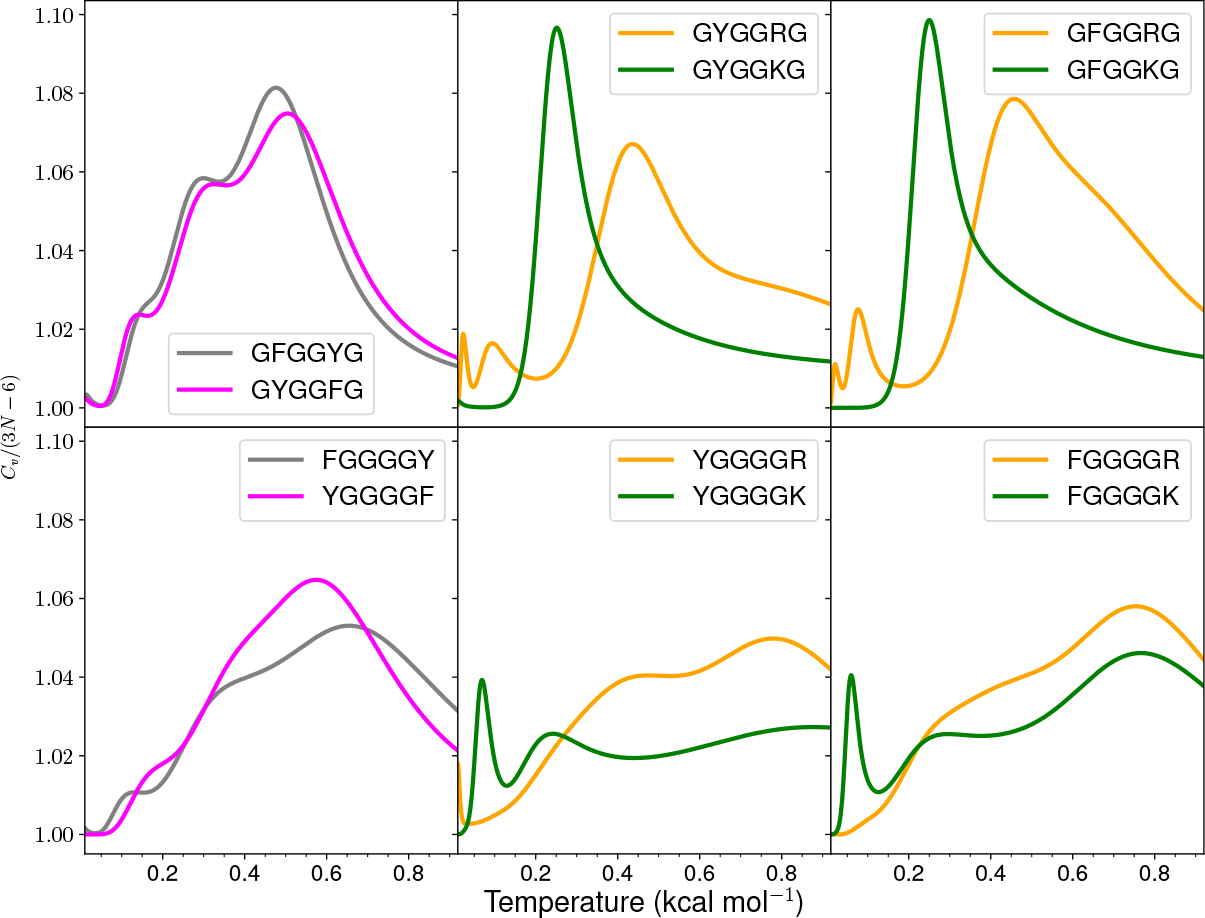
Heat capacity versus *k*_B_*T* for various hexapeptides.

**FIG. 8:**
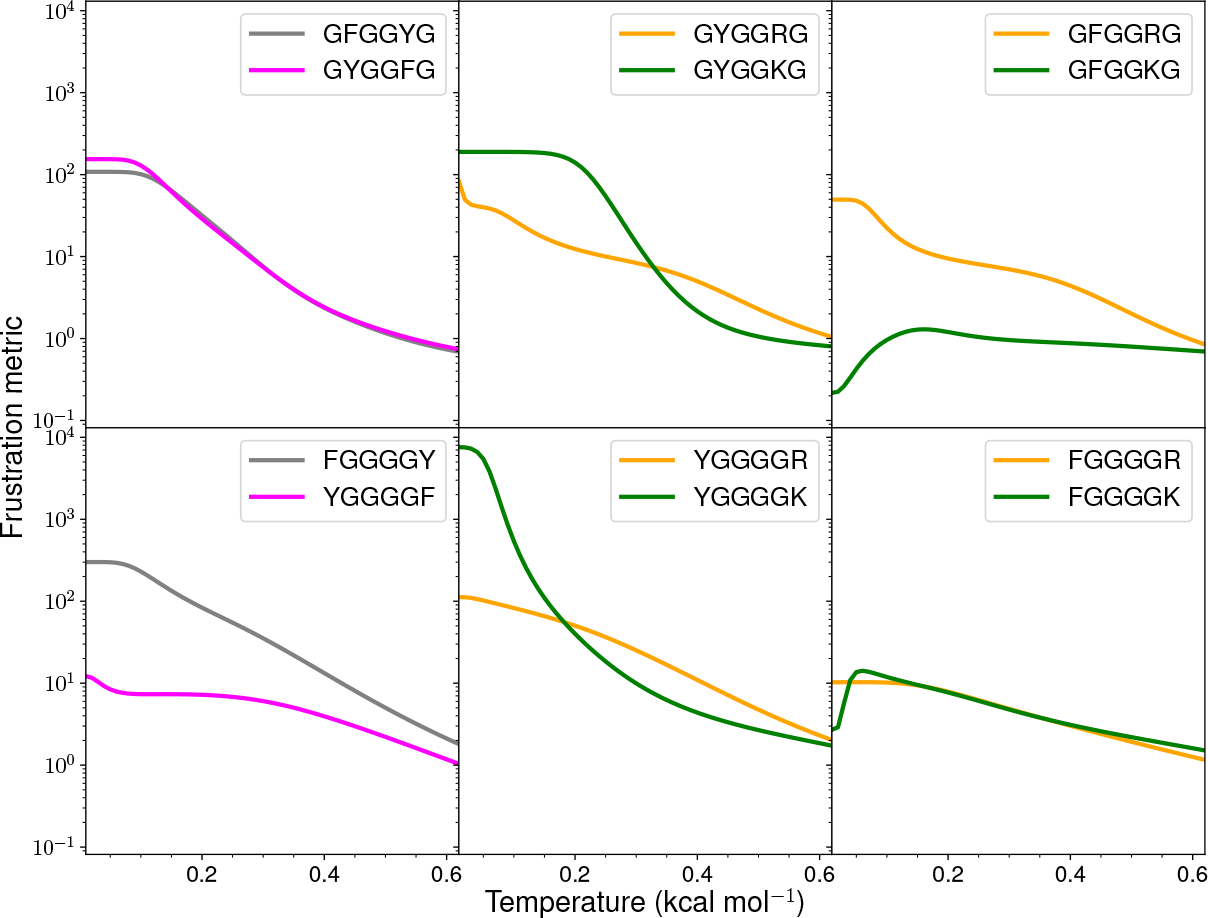
Frustration metric [*f*^r^(*T*)] versus *k*_B_*T* for various hexapeptides.

**FIG. 9:**
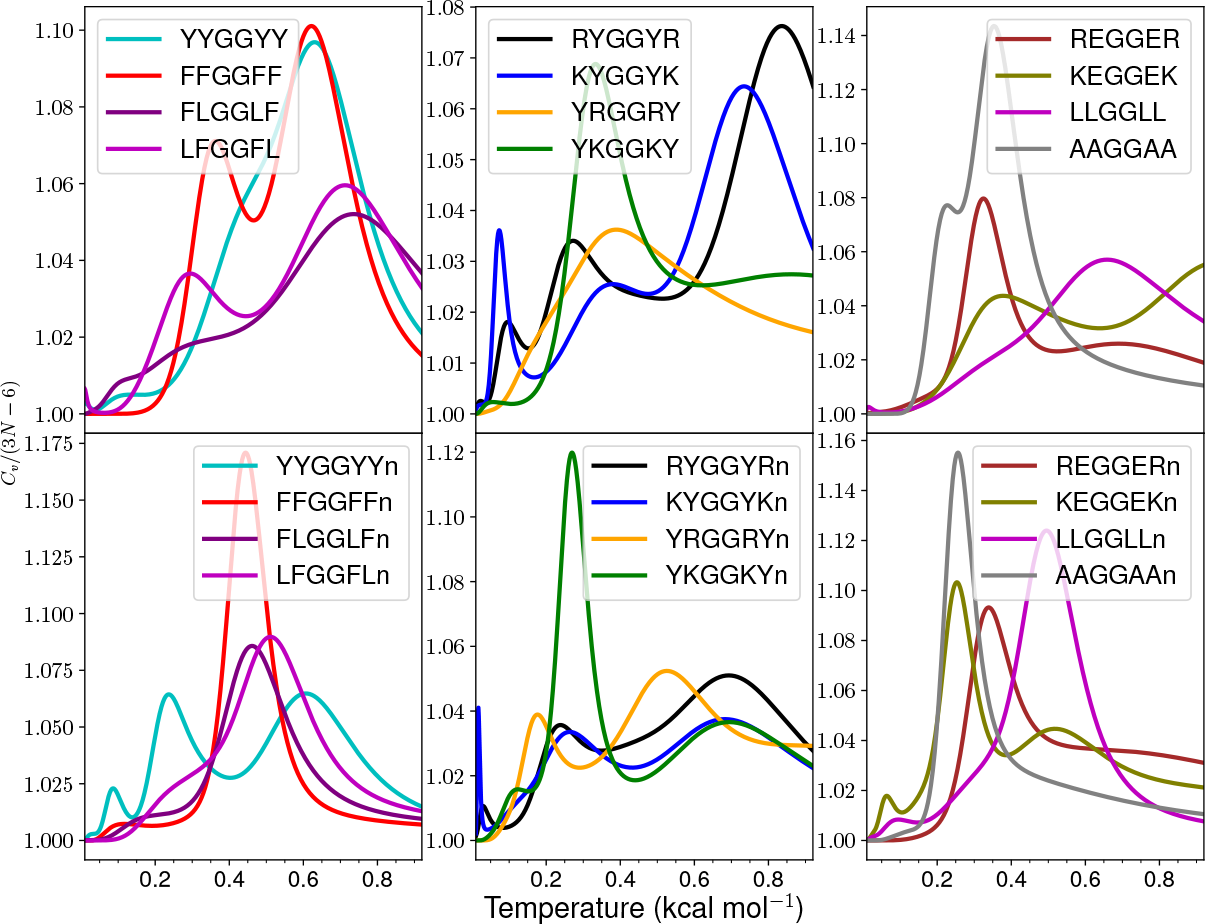
Heat capacity versus *k*_B_*T* for various hexapeptides. The uncapped peptide sequences end with a lowercase ‘n’.

**FIG. 10:**
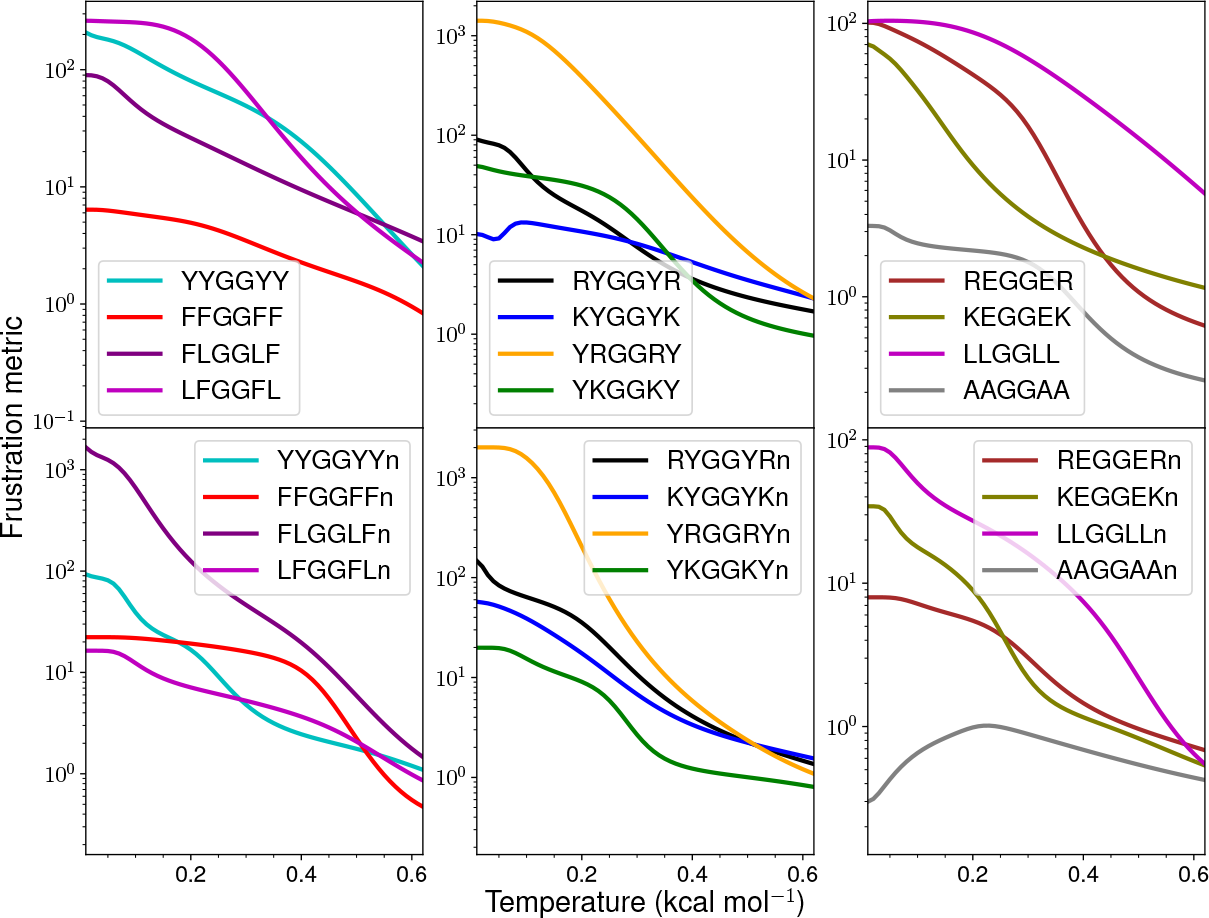
Frustration metric [*f* (*T*)] versus *k*_B_*T* for various hexapeptides. The uncapped peptide sequences end with a lowercase ‘n’.

**FIG. 11:**
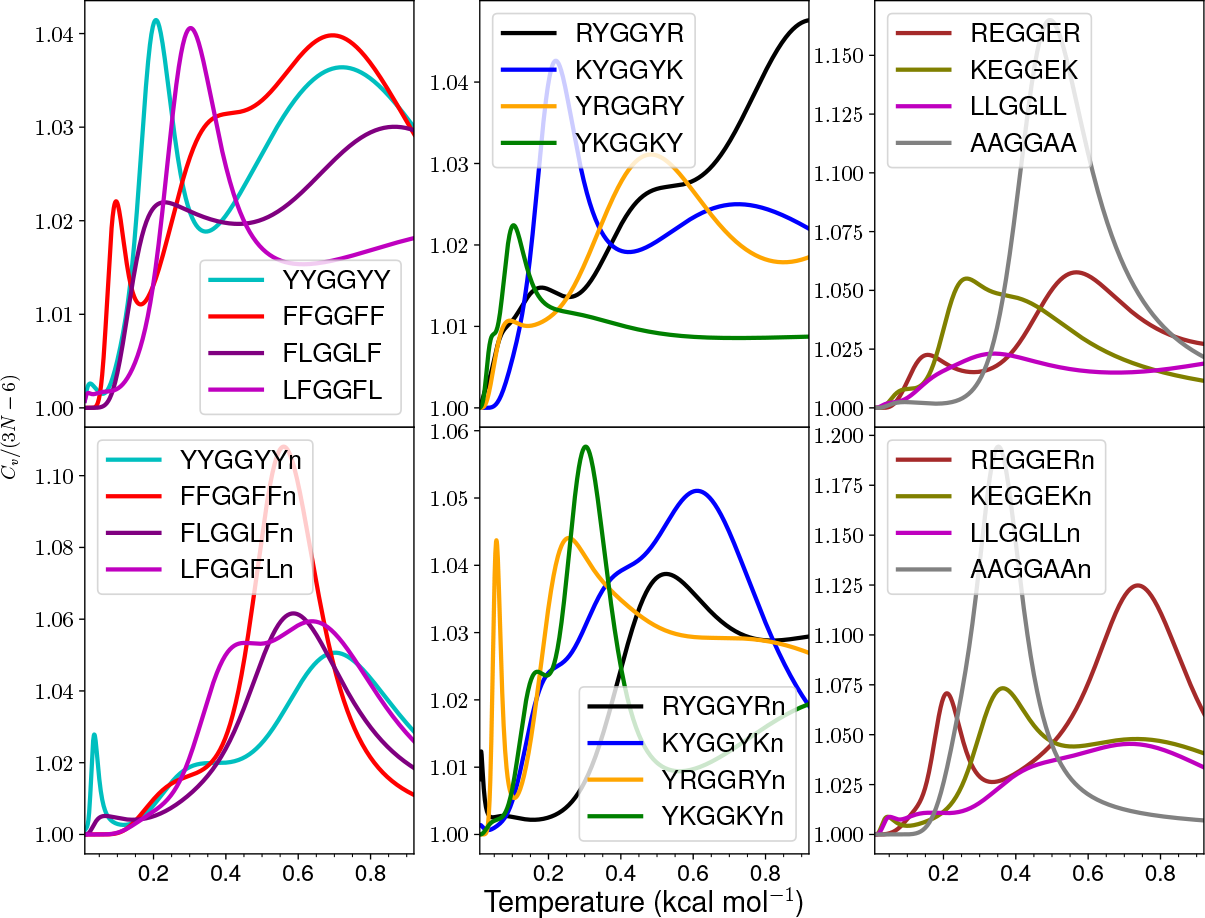
Heat capacity versus *k*_B_*T* for various hexapeptides modelled using FF19SB force field in AMBER20. The uncapped peptide sequences end with a lowercase ‘n’.

**FIG. 12:**
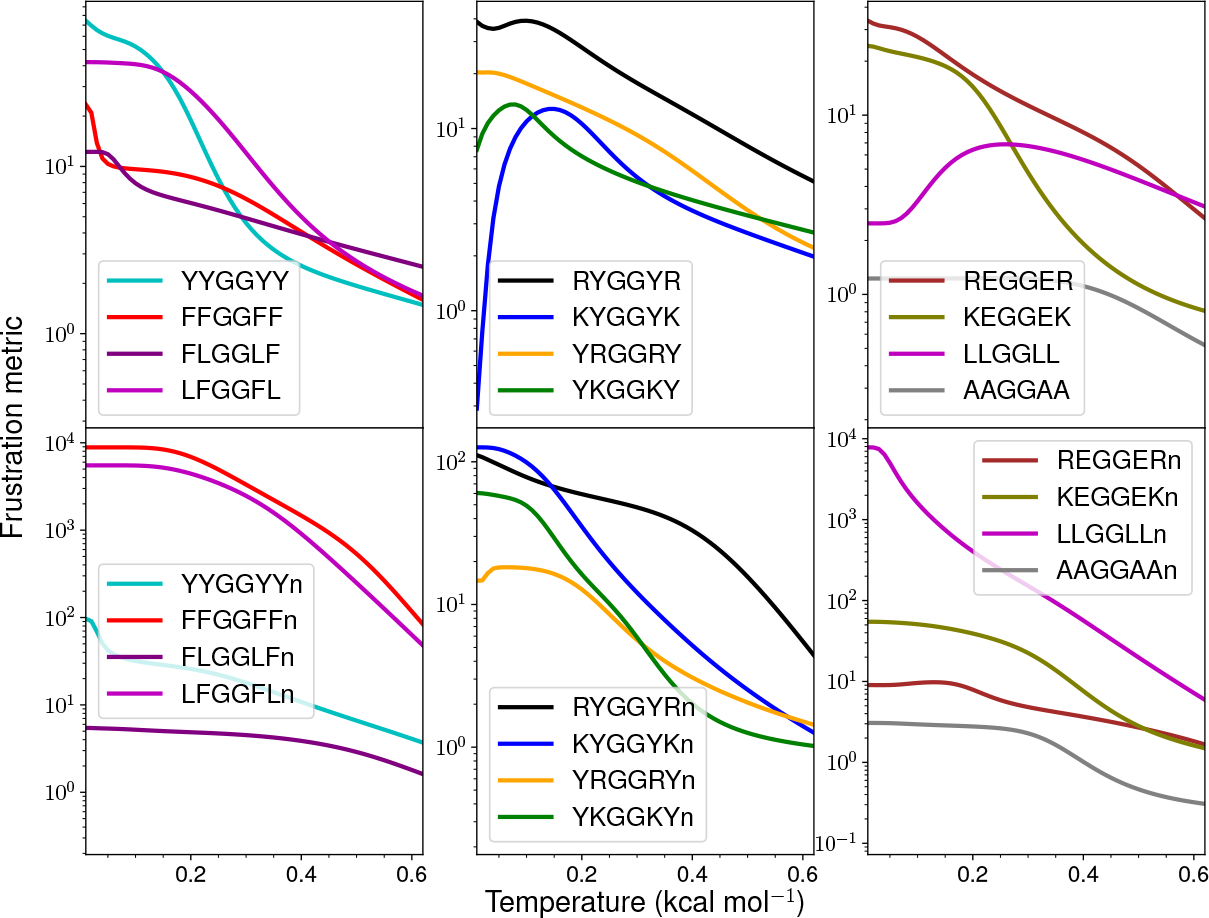
Frustration metric [*f* (*T*)] versus *k*_B_*T* for various hexapeptides modelled using FF19SB force field in AMBER20. The uncapped peptide sequences end with a lowercase ‘n’.

**TABLE I:** *k*_B_*T* in kcal mol*^−^*^1^ at which peaks and/or distinct inflection points are observed for selected peptides. The inflection points are marked with an asterisk (*^∗^*) and one very small peak is marked with ‘*i*’ (*^i^*).

| Hexapeptide | Peak/Inflection(*) temperatures |  |  |  |
| --- | --- | --- | --- | --- |
| GFGGFG | 0.158* | 0.323* | 0.493 | - |
| GYGGYG | 0.029 | 0.143 | 0.349 | - |
| GRGGYG | 0.028 | 0.053 | 0.495 | - |
| GKGGYG | 0.045 | 0.203 | 0.373 | - |
| GRGGFG | 0.022 | 0.450 | - | - |
| GKGGFG | 0.232 | 0.542 | - | - |
| FGGGGF | 0.098 | 0.415 | 0.600 | - |
| YGGGGY | 0.061* | 0.194* | 0.565 |  |
| RGGGGY | 0.120* | 0.474 | 0.676 | - |
| KGGGGY | 0.011 | 0.107 | 0.322 | 0.712 |
| RGGGGF | 0.128* | 0.462 | - | - |
| KGGGGF | 0.105 | 0.252 | 0.625 | - |
| FFGGFF | 0.363 | 0.622 | - | - |
| YYGGYY | 0.012 | 0.133 | 0.464* | 0.630 |
| FLGGLF | 0.131* | 0.348* | 0.736 | - |
| RYGGYR | 0.023<sup>i</sup> | 0.096 | 0.273 | 0.836 |
| KYGGYK | 0.073 | 0.380 | 0.734 | - |
| YKGGKY | 0.057 | 0.333 | 0.862 | - |
| REGGER | 0.325 | 0.690 | - | - |
| KEGGEK | 0.378 | 0.962 | - | - |
| AAGGAA | 0.227 | 0.354 | - | - |

**TABLE II:** *k*_B_*T* at which peaks and/or distinct inflection points are observed for selected hexapeptides. The inflection points are marked with an asterisk (*^∗^*) and a very small peak is marked with ‘*i*’ (*^i^*). The uncapped pe_3_p_3_tide sequences end with a lowercase ‘n’

| Hexapeptide | Peak/Inflection(*) temperatures |  |  |  |
| --- | --- | --- | --- | --- |
| GFGGYG | 0.014 | 0.165* | 0.299 | 0.477 |
| GYGGFG | 0.012 | 0.142 | 0.325 | 0.505 |
| GYGGRG | 0.023 | 0.094 | 0.436 | - |
| GYGGKG | 0.250 | - | - | - |
| GFGGRG | 0.020 | 0.076 | 0.457 | - |
| GFGGKG | 0.250 | - | - | - |
| FGGGGY | 0.132 | 0.430* | 0.622 | - |
| YGGGGF | 0.197* | 0.574 | - | - |
| YGGGGR | 0.475 | 0.779 | - | - |
| YGGGGK | 0.069 | 0.242 | 0.743 | - |
| FGGGGR | 0.468* | 0.754 | - | - |
| FGGGGK | 0.060 | 0.296 | 0.766 | - |
| LFGGFL | 0.293 | 0.703 | - | - |
| YRGGRY | 0.386 | - | - | - |
| LLGGLL | 0.015<sup>i</sup> | 0.355* | 0.659 | - |
| FFGGFFn | 0.117 | 0.444 | - | - |
| YYGGYYn | 0.033* | 0.086 | 0.237 | 0.607 |
| LFGGFLn | 0.287* | 0.511 | - | - |
| FLGGLFn | 0.208* | 0.461 | - | - |
| RYGGYRn | 0.030 | 0.239 | 0.693 | - |
| KYGGYKn | 0.017 | 0.264 | 0.682 | - |
| YRGGRYn | 0.175 | 0.518 | 0.876* | 0.918 |
| YKGGKYn | 0.123 | 0.270 | 0.694 | - |
| REGGERn | 0.339 | - | - | - |
| KEGGEKn | 0.064 | 0.253 | 0.518 | - |
| LLGGLLn | 0.095 | 0.470 | - | - |
| AAGGAAn | 0.256 | - | - | - |

**TABLE III:**
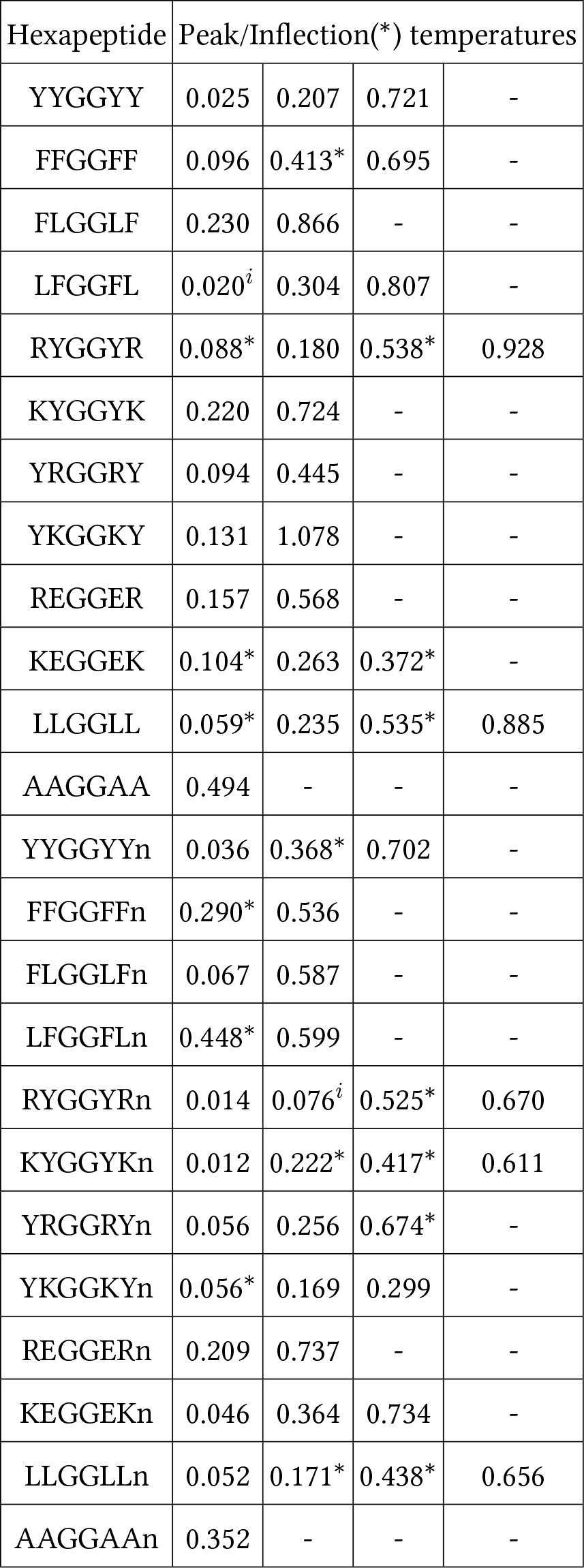
*k*_B_*T* at which peaks and/or distinct inflection points are observed for selected hexapeptides using the FF19SB force field in AMBER20. The inflection points are marked with an asterisk (*^∗^*) and two very small peaks are marked with ‘*i*’ (*^i^*). The uncapped peptide sequences end with a lowercase ‘n’.

**TABLE IV:**
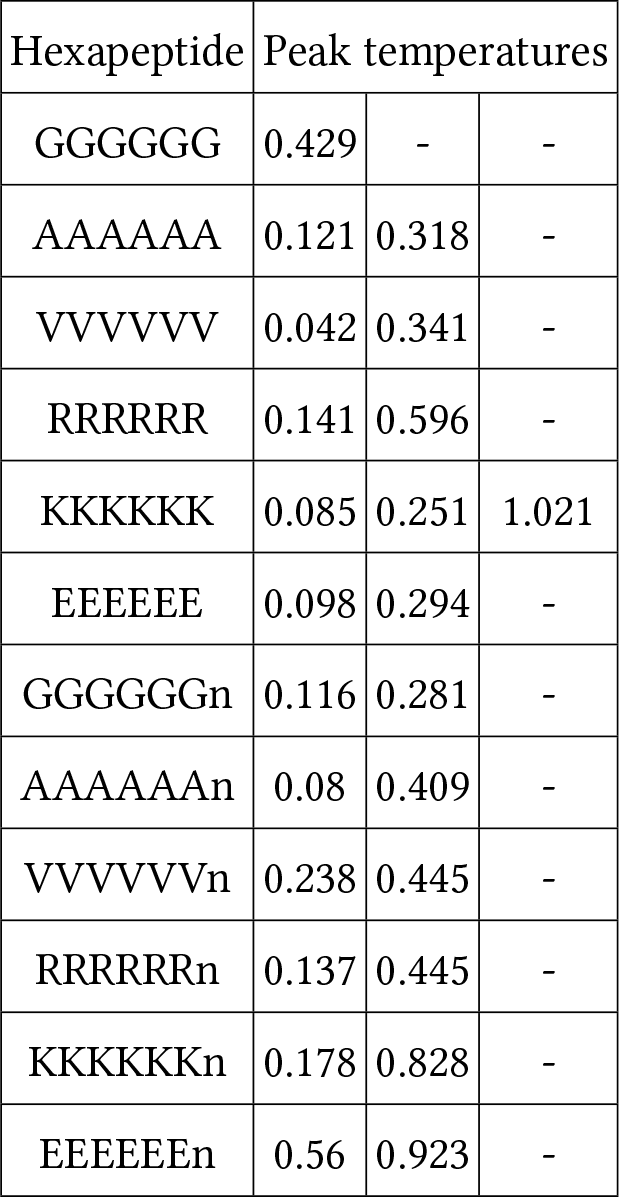
*k*_B_*T* at which peaks and/or distinct inflection points are observed for selected hexapeptides using the FF99IDPS force field. The uncapped peptide sequences end with a lowercase ‘n’.

## Computational protocol

The AMBER input files can be created by following the method given at https://wikis.ch.cam. ac.uk/ro-walesdocs/wiki/index.php/Preparing input files for a peptide using AMBER. The steps to clone the softwarewales repository are given at https://wikis.ch.cam.ac.uk/ro-walesdocs/wiki/ index.php/Git Workflow. The steps to obtain the A12GMIN, A12OPTIM, and PATHSAMPLE executables are given at https://wikis.ch.cam.ac.uk/ro-walesdocs/wiki/index.php/Compiling Wales Group codes using cmake. The example input files for running basin-hopping parallel tempering and discrete path sampling are given in the next section. The explanation of all the keywords can be found at https://www-wales.ch.cam.ac.uk/GMIN.doc/node7.html for the data file, https://www-wales.ch.cam.ac.uk/OPTIM.doc/node4.html for the odata.connect and odata.addminxyz files, and https://www-wales.ch.cam.ac.uk/PATHSAMPLE.2.1.doc/node6.html for the pathdata file. The explanation of how to add the minima obtained using A12GMIN to a PATHSAMPLE database is given at https://wikis.ch.cam.ac.uk/ro-walesdocs/wiki/index.php/ Adding several minima obtained using GMIN (maybe using BHPT) to min.data. The creation of input files and executables required for heat capacity analysis is given at https://wikis.ch.cam.ac.uk/ro-walesdocs/wiki/index.php/ Decoding heat capacity curves. The disconnectivity graph can be constructed using the disconnectionDPS program documented at https://www-wales.ch. cam.ac.uk/disconnectionDPS.doc/. Example input files used for one of the peptides are given below.

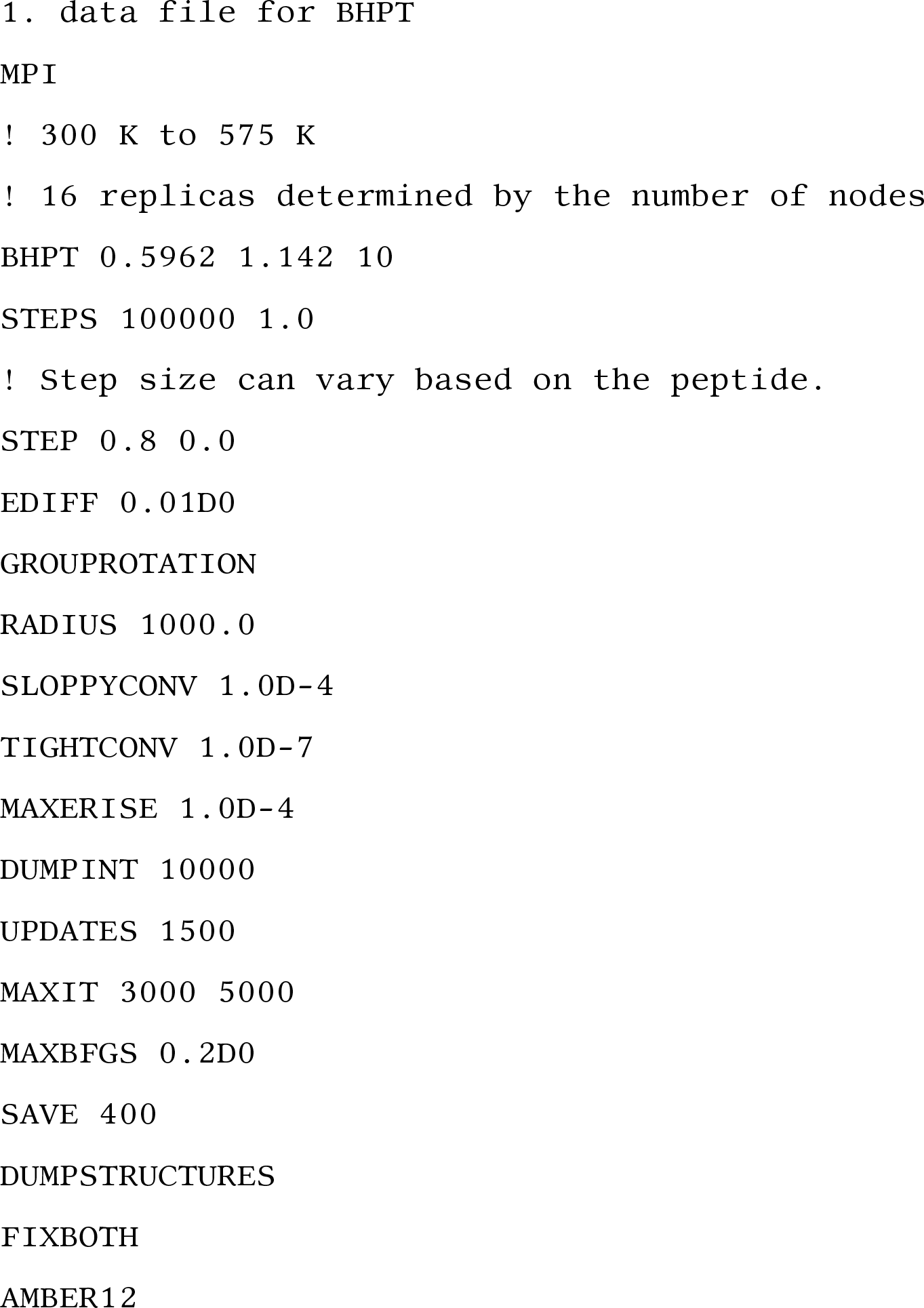
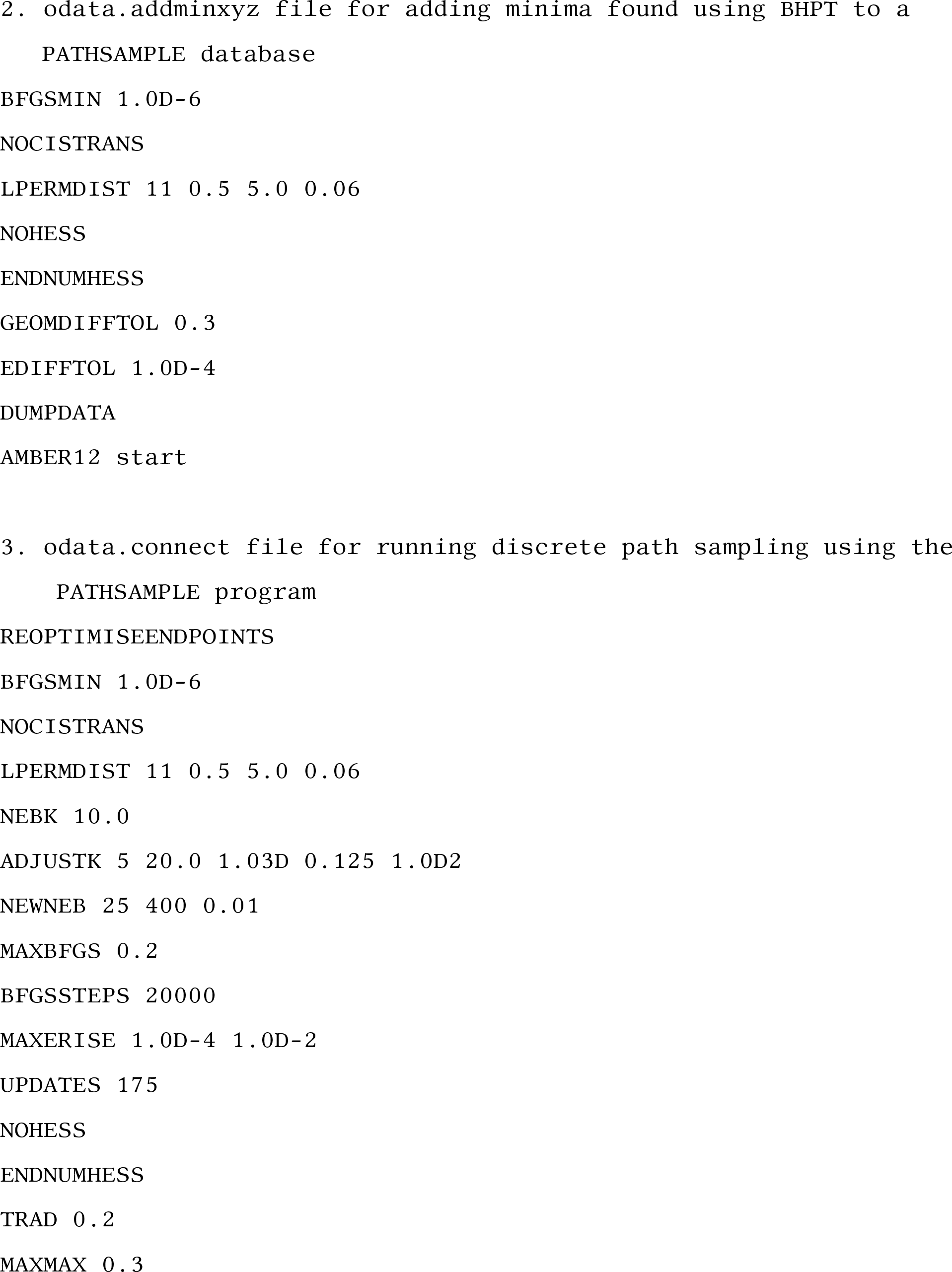
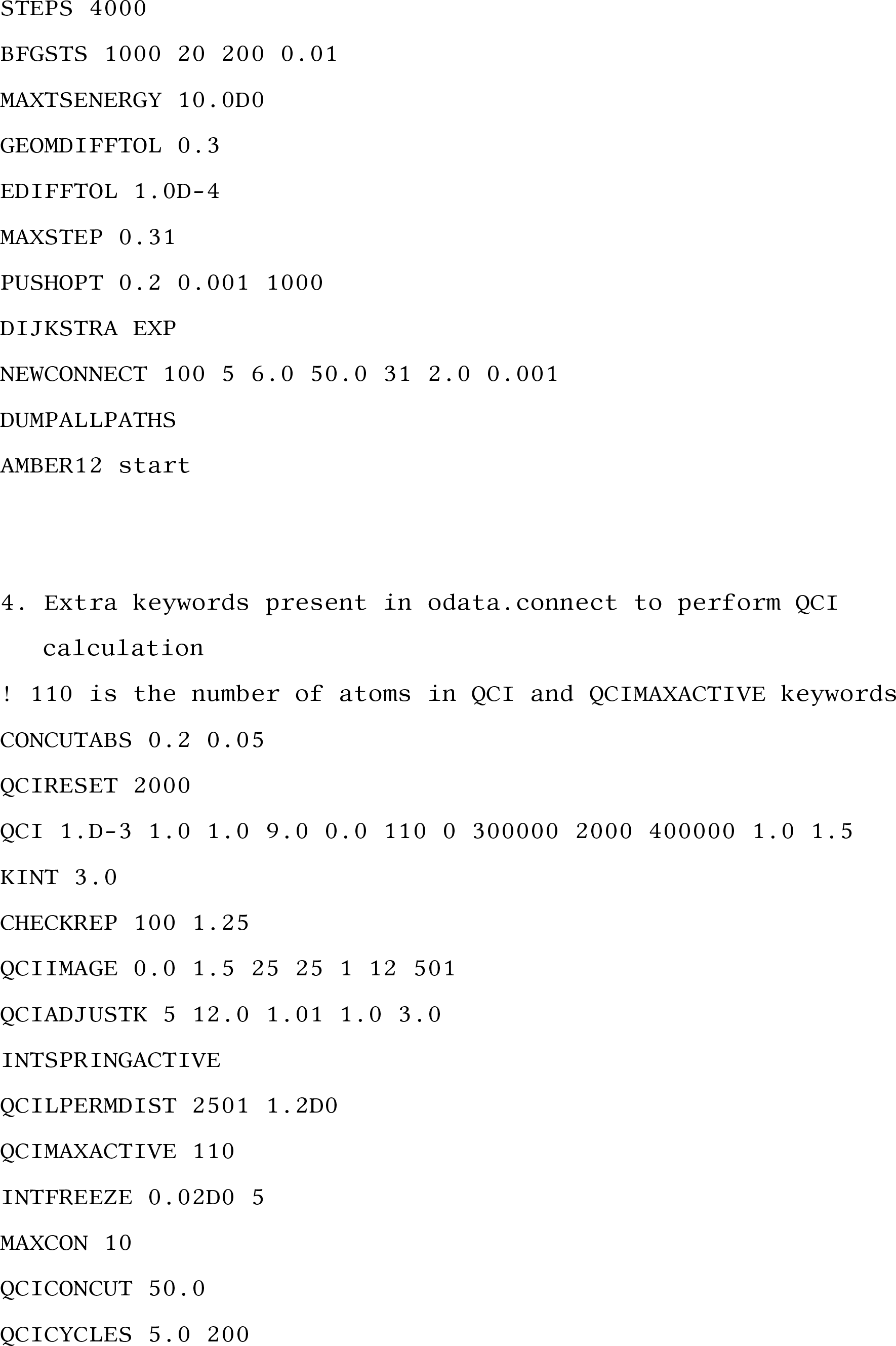
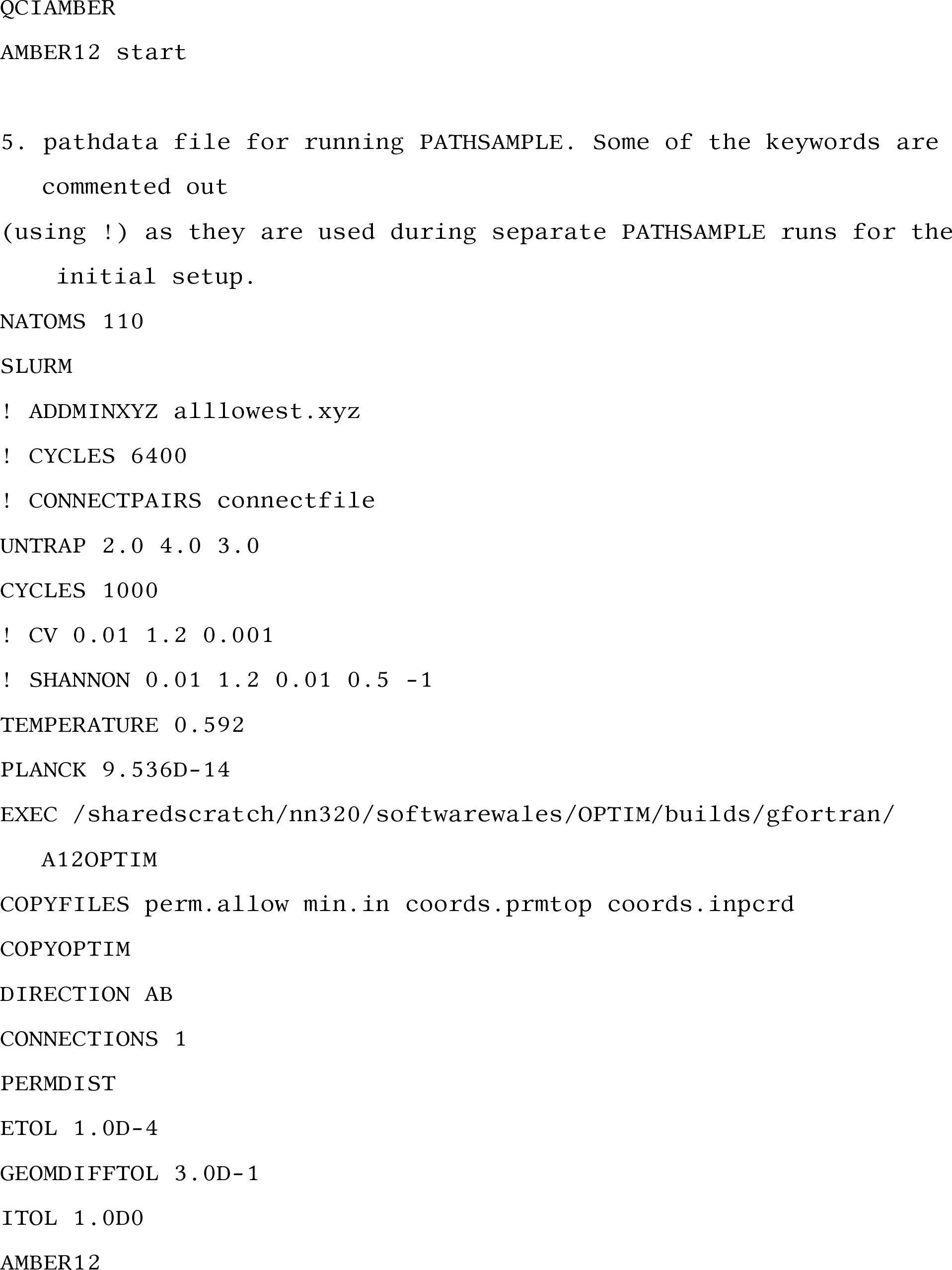
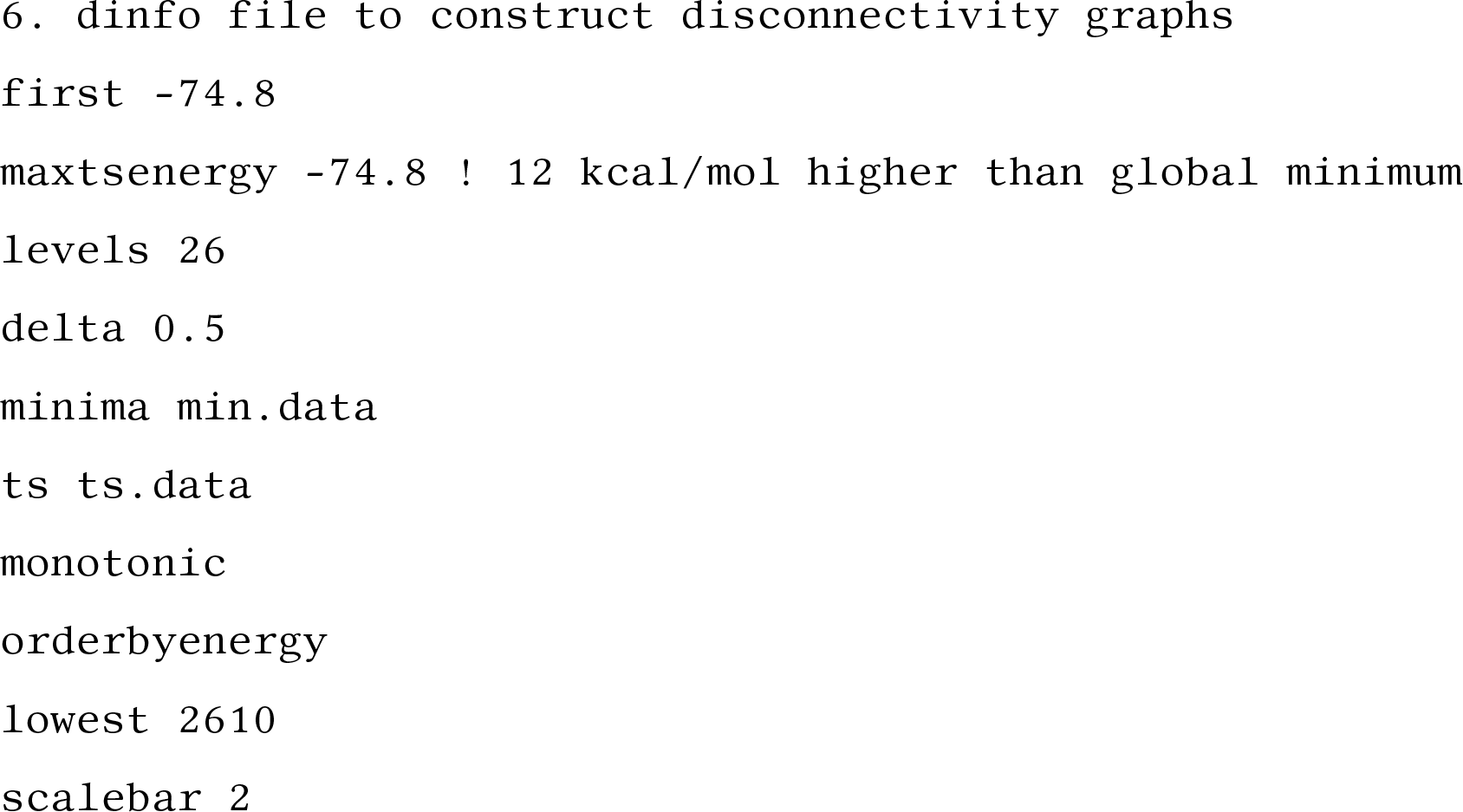

